# Binning-based Methods for Model Fitting and Data Splitting Improve Machine Learning Imbalanced Regression

**DOI:** 10.1101/2025.06.26.661884

**Authors:** Husam Abdulnabi, J. Timothy Westwood

## Abstract

Modelling quantitative genomics data remains challenging due to imbalanced target distributions, non-independent observations, and substantial computational costs. This study examines the challenges and current methods, proposes new methods, applies these methods across two case studies: a controlled setting with synthetically generated data and on real genomics datasets.

Effective modelling of imbalanced data starts with density estimation. Density estimation via binning facilitates downstream analytical steps. Minimum Transformation Binning is a novel method that improves current binning methods for density estimation.

Data splitting of imbalanced and non-independent observations is best accomplished by objective data splitting which entails randomly generating splits, scoring them, and selecting the best scoring one. This study conducts an empirical comparison of scoring functions toward an optimal objective data splitting method, finding the use of all-against-one absolute relative difference to be the best scoring function.

Model fitting methods can be divided into loss functions and data balancing. Two methods have been previously proposed based on weighting error functions by the reciprocal or complement of the density estimation. This study conducts a comparison of these methods on the case studies, finding that reciprocal density weighting significantly outperforms non-weighted and complement density weighting.

This study also tests model fitting with reciprocal weighted sampling per epoch for imbalanced regression. While this method does not generally improve performance, it uses significantly less resources in model fitting which can be beneficial for hyperparameter optimization.

Together, this study provides a unified framework for effective modelling of imbalanced regression, especially imbalanced genomics data.

## 1 Introduction

Effective machine learning models for genomic data can improve future experimental design and reveal biological concepts not otherwise accessible. This study defines genomic modelling as sequence-to-function, where the inputs are sequences (primarily nucleotide sequences) and the outputs (i.e., targets) are experimentally measured values associated with those input sequences. While the experimentally measured values can be binarized and modelled as a classification task, the values often contain rich quantitative information (Belgacem & Delaplace, 2025). Thus, this study focuses on the regression of genomic data. Despite advances in data quality and model architectures, modelling genomic data remains challenging due to target value imbalance, non-independence, and large resource requirements.

First, many genomic assays, such as chromatin immunoprecipitation (ChIP) experiments, often produce right-skewed values where higher values are increasingly rare. Targets that exhibit uneven density across their range are known as imbalanced. Target imbalance complicates model scoring, data splitting, and model fitting. Modelling imbalanced continuous targets (i.e., imbalanced regression) often benefits from the accurate estimation of the density of the target distribution. However, density estimation is not straightforward for imbalanced distributions, especially highly skewed distributions.

Second, modelling genomic data entails partitioning sequences into windows that may overlap or interact, rendering them non-independent. Together with imbalanced targets, genomic data requires special care in data splitting.

Third, effective modelling of genomic data requires large datasets and increasingly complex machine learning models that require substantial computing resources. Although advances in hardware and software have reduced some barriers, efficient modelling remains crucial for accessibility, reproducibility, and optimization.

This study examines the challenges and current solutions to density estimation (**Section 1.1**), data splitting (**Section 1.2**), and model fitting (**Section 1.3**). This study proposes novel binning-based methods to improve the modelling of imbalanced genomic regression tasks, with applicability extending well beyond genomics, described in more detail in **Section 2**. Finally, we evaluate existing and proposed methods in a controlled environment with synthetically generated data in **Section 3** and on real-world genomics datasets in **Section 4**.

### 1.1 Density Estimation

Kernel Density Estimation (KDE) is a widely used approach to estimate density (Scott, 2015; Silverman, 1986). This study, however, focuses on non-overlapping binning methods as they facilitate downstream tasks such as data splitting and are also easier to interpret. Representing the density of a distribution using binning yields a *B*-dimensional vector: let **p** = (*p*_1_, … *p_B_*) ∈ ℝ*^B^*, where each dimension corresponds to a specific bin and the values are bin densities (**Eq. 1**).

Eq. 1. **Bin Density**

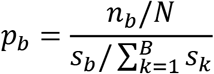

Where:

*p_b_* = density of bin *b*
*n_b_* = number of members belonging to bin *b*
*N* = number of members
*s_b_* = width of bin *b*
*B* = number of bins

There are three major considerations for effective binning. First, the number of bins; too few bins obscure the underlying distribution structure, while an excessive number becomes impractical for downstream applications. Second, the bin widths; excessively large bins lose specificity. Third, the minimum number of observations per bin; sparse bins complicate data splitting and yield unstable local performance measurements.

Histogram binning partitions the target range into equal-width bins. This approach often fails in sparse regions by yielding bins with too few observations. Alternatively, bins with variable widths can be used to ensure a minimum number of values per bin. Notably, the widths of the bins need to be considered for density estimation. For example, observations belonging to a bin *A* with twice the number of values and twice the width of another bin *B* have the same density as those belonging to bin *B*.

Quantile and rank binning are two variable-width approaches that yield bins with an approximately equal number of values in each bin. While the minimum number of observations can be used to determine bin boundaries, this often results in a trade-off: enforcing a small minimum (e.g., 20 observations) in a large dataset (e.g., 100,000 observations) yields an unmanageable number of bins (e.g. 5,000 bins), while a larger minimum may produce bins that lack necessary resolution. More sophisticated variable-width methods, such as Bayesian Blocks, determine bin boundaries by identifying significant changes in density through the optimization of a fitness function (Scargle et al., 2012). This approach, however, is computationally taxing (approaching O(*N*^2^) complexity) and non-parallelizable, which is concerning for genomic datasets that often contain hundreds of thousands of observations. Furthermore, these methods can yield bin widths that are less interpretable, as neighboring bins may fluctuate in size without a monotonic trend.

For unimodal tailed distributions—including right-skewed, left-skewed, and bell-shaped distributions—an alternative is to apply a monotonic transformation that compresses the tails, perform histogram binning in the transformed space, and map the bins back to the original scale. This study refers to this strategy as Transformation binning. After transformation, the number of bins can be selected as the maximum required to satisfy a desired minimum count per bin. Transformation binning is typically applied to data shifted so that the mode is centered at 0. This approach yields interpretable bin widths that monotonically increase as they move away from the mode. However, different parameters of the monotonic transformation yield different bin configurations, and it remains unclear which parameterization produces the most desirable result. While a log transformation is a common choice for right-skewed data, its fixed functional form may yield suboptimal widths or counts when mapped back to the original scale.

The Box-Cox (Box & Cox, 1964) and Yeo-Johnson (Yeo & Johnson, 2000) transformations are objective power transformations that aim to make distributions more symmetric and bell-shaped. However, using these transformations for transformation binning may still produce bins with undesirable widths: selecting transformation parameters that satisfy specific bin properties remains non-trivial.

This study proposes Minimum Transformation Binning: a novel objective transformation method that optimizes the monotonic transformation for a desired minimum number of values per bin and maximum bin widths, while automatically determining the optimal number of bins (detailed in **Section 2.1**). It is also computationally efficient and parallelizable.

### 1.2 Data Splitting

Machine learning modelling entails dividing the data into at least a training and a testing set. The training set is used to fit the model, and the testing set is used to assess (i.e. score) the model. The data should be divided such that each set contains observations that are independent. Non-independent observations that are spread across different sets affect both model fitting and scoring which leads to improper model evaluation, known as data leakage. To prevent this, non-independent observations need to be grouped together prior to splitting.

Moreover, the data should be divided such that the outputs (i.e. target values) of each set are identically distributed, a property determined here through binning. Notably, there exist data splitting methods that also consider or solely consider the inputs (i.e. features) such as SPlit (Joseph & Vakayil, 2022). This study, however, focuses on data splitting solely on the output values. In the context of sequence-to-function modelling, determining or extracting definitive features from the raw sequence inputs is not trivial, making output-based partitioning a more robust and direct approach.

Furthermore, each set must contain a sufficient number of observations per bin to ensure robust model fitting and scoring. Maintaining identically distributed sets ensures that target imbalance is represented proportionally across all partitions. This is particularly critical for under-represented, low-density values. Additionally, the split must achieve the desired proportions of observations (e.g. a 70% training and 30% testing split).

Observations from genomics data are often windows of larger reference sequences, typically chromosomes. These windows can be overlapping and thus must be grouped (**Fig. 1**). Depending on the experimental assay that produced the target values, there must also be consideration for grouping adjacent non-overlapping windows as they may interact (Kaul et al., 2020). In tasks involving multiple reference sequences, data can be partitioned by assigning entire reference sequences to specific sets, thereby confining overlapping and adjacent windows. However, because reference sequences vary in size and window density, this approach often fails to achieve the desired split proportions. Alternatively, as is used in this study, neighboring observations can be grouped based on a determined distance threshold.

**Fig. 1.**
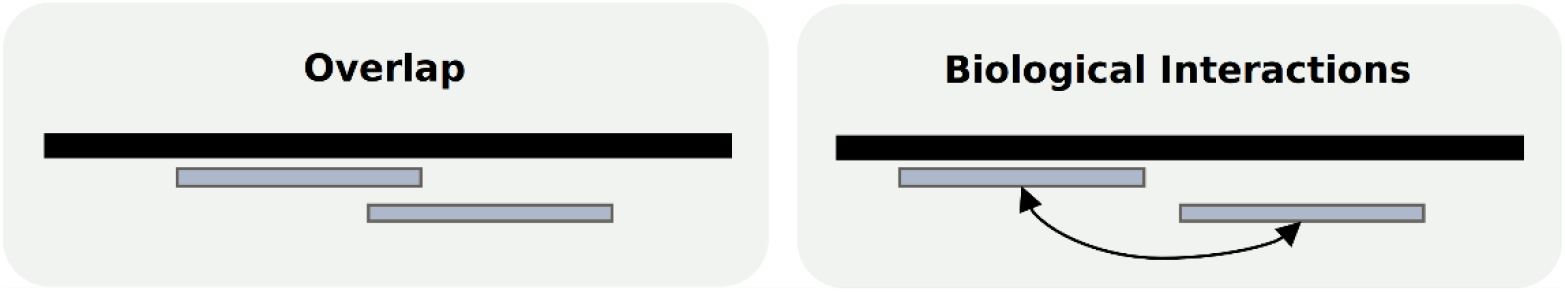
Windows for modelling can be non-independent by overlapping or having biological interactions. Black lines represent reference sequences and blue lines represent windows of the reference sequences.

Data can be split by random sampling groups without replacement, known as random split. This method accounts for the desired proportions of each set and guarantees independent sets but does not guarantee identically distributed data. One method to guarantee identically distributed data is stratified split that performs proportional random sampling without replacement of each stratum (i.e., each bin). Still, stratified split is not compatible with non-independent observations as it cannot be applied to groups containing members belonging to different bins.

For identically distributed sets of non-independent observations, one approach entails generating multiple candidate data splits through randomly splitting groups (which ensures independent sets) and then selecting the split that has the most identically distributed sets. The method of scoring splits and selecting the best one is sometimes referred to as “Best of N”. Specifically using this method on non-independent splits is referred to hereafter as objective splitting. More involved optimization algorithms (e.g. genetic algorithms, simulated annealing) can be used, but this study focuses on using random optimization for its simplicity.

For each candidate split and for each target value, the distribution of the values of each set (and optionally of all the observations) is represented as a vector. In this study, distributions are represented by their bin densities and the same bins are used on different distributions, yielding comparable same-length vectors. The similarity of two vectors can be determined by metrics such as the Kullback-Leibler (KL) divergence. When this shared binning scheme has variable bin widths, metrics are weighted by bin size so that larger bins contribute proportionally more to the score.

The vectors can be compared to the corresponding vector of all observations, referred to as an all-against-one comparison (**Eq. 2**). Alternatively, the vectors of the sets can be compared to each other pairwise, referred to as an all-against-all comparison (**Eq. 3**). When there are multiple comparisons, the comparisons can be summarized as the mean of the comparisons.

All-against-one scoring for data splitting has been explored before: using equal-sized bins with the KL divergence or Wasserstein distance (Jami et al., 2025). Comparisons of vectors with unequal-sized bins and pairwise comparisons for objective data splitting have rarely if ever been done but we do not claim novelty.

Objective splitting with different metrics can yield different data splits as different metrics emphasize different aspects of distributional similarity. To the best of our knowledge, there is no empirical study comparing metrics for objective data splitting. **Section 3** is a case study that includes experimenting with different scoring functions for objective data splitting.

Eq. 2. **All-against-one comparison**

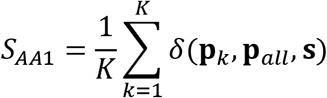

Where:

*K* = number of sets
**p**_all_ = bin density of all observations
**s** = (*s*_1_, …, *s_B_*) ∈ ℝ*^B^*
*δ* = weighted similarity metric

Eq. 3. **All-against-all comparison**

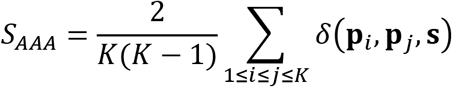

### 1.3 Model Fitting

In some ML tasks, some values are more important than others. In other words, the importance, often called relevance, of values is not uniform. For example, in fraud detection, extreme values have more importance than values closer to the mean. Weights are assigned to values based on their importance, referred to here as relevance weights, and are used in metrics. For regression tasks, this is known as utility-based regression and can involve the binarization of continuous values (Torgo et al., 2013). In this study, however, all ranges are equally important.

This study defines genomics modelling as sequence-to-function tasks. Neural Networks (NNs) have been proven effective for such tasks and thus are the focus of this study. There are two categories for model fitting methods for imbalanced data: loss function and data balancing.

#### 1.3.1 Loss Function

Loss Function methods assign higher weights to observations in less-dense regions to give them more influence than observations in more-dense regions. The Balanced Mean Squared Error (Balanced MSE) weights the observations by the inverse of the density estimation (Ren et al., 2022). Conversely, DenseWeights weights the observations by the complement of the density estimation (Steininger et al., 2021). For better interpretability, we refer to balanced MSE and MSE weighted by DenseWeights as reciprocal-weighted MSE (recip-MSE) and complement-weighted MSE (comp-MSE), respectively. To the best of our knowledge, an empirical comparison of these two methods has not been performed previously. The two loss functions are tried in a controlled setting in **Section 3** and on real genomics datasets in **Section 4**.

#### 1.3.2 Data Balancing

Data balancing methods aim to reduce data imbalance used to fit the model by sampling and/or synthetically generating observations.

A straightforward sampling method is reciprocal-weighted random sampling wherein observations are weighted by the inverse of the density estimation. Sampling can be performed without replacement or with replacement. Sampling without replacement prunes over-represented data and results in smaller amounts of data to fit the model which may require less computing resources. The proportion of sampling is not clear, but data can become more balanced with smaller sampling proportions. Sampling without replacement is especially effective when there is redundant over-represented data as it is expected to yield at least similar performance while reducing computational costs.

Sampling with replacement often results in replicating less-represented observations, pruning more-represented observations, and balanced data. Again, the proportion of sampling is not clear, but proportions less than 1 result in smaller amounts of data which may reduce computational costs.

There exist more involved sampling methods that consider the observations’ inputs. For example, Prototype Selection methods cluster the observations by their inputs and select maximally diverse representatives from each cluster, yielding a compact yet informative subset of data (Garcia et al., 2012). Similarly, Synthetic Minority Oversampling Technique for Regression with Gaussian Noise (SMOGN) augments less-represented data to generate additional less-represented data (Branco et al., 2017). These methods, however, are not straightforward for sequence inputs and thus are not tried in this study.

Repeated data balancing prior to model fitting can yield different datasets which may yield substantially different models. NNs provide a unique opportunity for data balancing as they are fit in cycles called epochs. Instead of data balancing before model fitting, data balancing can occur prior to each epoch. Weighted random sampling at each epoch is referred to here as weighted epoch sampling. For imbalanced classification, weighted epoch sampling with reciprocal frequency weights has been applied before (Tomei et al., 2021). For imbalanced regression, however, it has yet to be investigated to the best of our knowledge. This method is evaluated with or without replacement and with different sampling proportions in a controlled setting in **Section 3** and on real genomics datasets in **Section 4**.

## 2 Novel Methods

### 2.1 Minimum Transformation Binning

Minimum Transformation Binning searches for a scale factor applied to the data before a monotonic transformation such that the resulting histogram satisfies two constraints: (1) each bin in the transformed space contains at least a minimum number of observations and (2) the corresponding bin widths in the original (untransformed) scale do not exceed a maximum allowed size (**Algorithm 1**). Among all feasible configurations, the procedure selects the maximum number of bins that still satisfies these constraints. In practice, the algorithm is parallelized over candidate bin counts and scale factors.

#### Algorithm 1.

Minimum Transformation Binning Inputs:

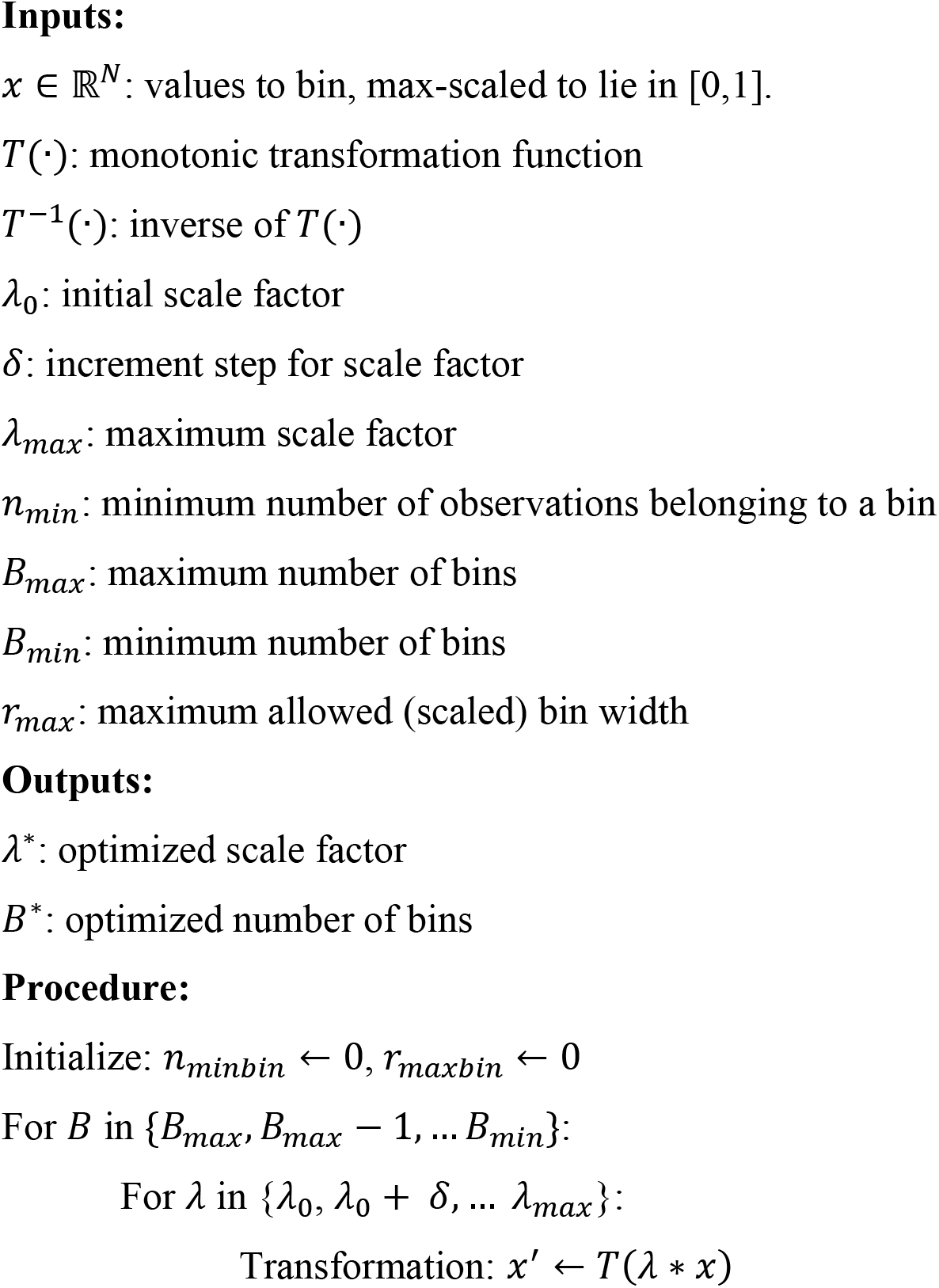
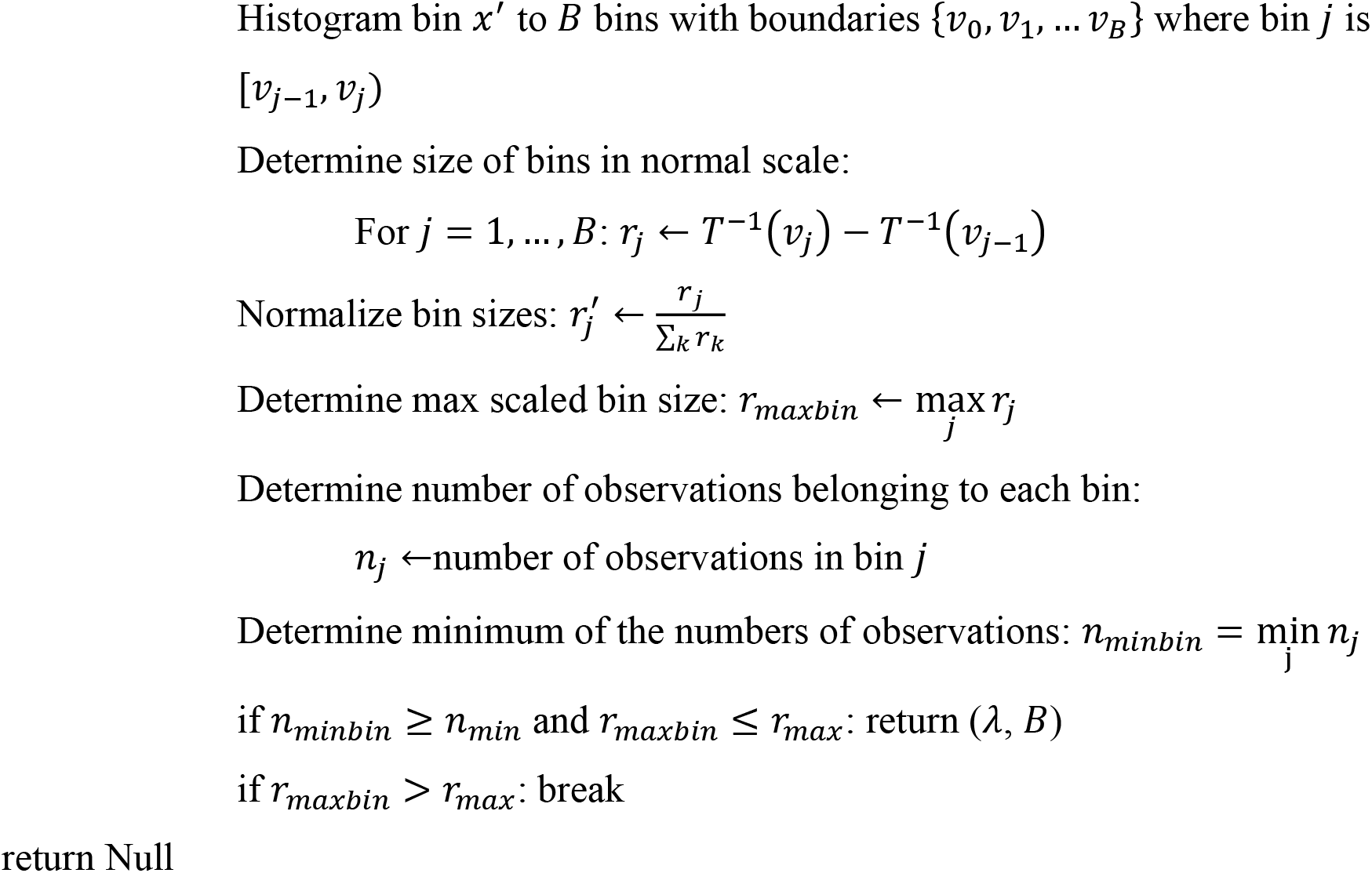

## 3 Case Study: Synthetic Right-Skewed Data

Synthetically generated data provides several advantages over real-world data for evaluating methods for imbalanced regression. First, the data attributes can be controlled, including the complexity of the data, the types and amounts of added noise, and degree of imbalance. Second, the same or very similar model architectures can be used to generate the data and later model the data. This obviates the need to search model configurations (discussed later). This also ensures that the data is able to be modelled and provides guardrails for data generation. Too little noise and/or imbalance yields data that are too easily modelled, whereas too much noise and/or imbalance yields data that is nearly impossible to sufficiently model. Third, the minimum number of observations to adequately model the data can be determined, ensuring effective computational cost. Fourth, coupled with the third point, data generation can be repeated at multiple steps in the process, ultimately yielding robust findings.

The following case study has two primary objectives. The first is to determine the optimal scoring function for objective data splitting. A total of 11 scoring metrics were tested including ones used before in other studies as well as ones that have not. Furthermore, all-against-one and all-against-all methods are compared. Together, this provides the first comprehensive test of objective data splitting methods, to the best of our knowledge. The second is to determine the efficacy of model fitting methods. Specifically, different loss functions and different Epoch sampling methods are tested toward improving the modelling of imbalanced data. While recip-MSE and comp-MSE have been explored before, they have yet to be directly compared to the best of our knowledge. Also, Epoch Sampling has not been formally tested for imbalanced regression, to the best of our knowledge. Together, these approaches provide a comprehensive test of model fitting methods for imbalanced data.

### 3.1 Methods

See https://github.com/Husam94/BinMeths/CS-Synth/ for complete details.

#### 3.1.1 Neural Networks

Inspired by Steininger et al., 2021, a NN with 3 Dense layers each with 10 nodes, ReLU activation functions between the Dense layers, and a single output node was used, referred to here as the Base NN. Batch normalization layers were added between the Dense layers as they are found in most current NN architectures.

#### 3.1.2 Data Generation

A 10-feature input was generated by sampling uniform distributions with a range [-1, 1]. A total of 1,000,000 10-feature inputs were generated. A Base NN was randomly initialized by Kaiming Initialization and produced corresponding outputs from the inputs. The pairs of input-output are referred to as original data. This procedure was repeated 20 times, generating 20 sets of original data from 20 different initializations of Base NNs, referred to as synthetic repeats. The outputs of the synthetic repeats were visually inspected for bell-shaped distributions. Most randomly initialized NNs in this study yielded bell-shaped output distributions, while a small fraction did not. These few are assumed to have poor initializations that are assumed to obscure the mapping of input to output by other NNs downstream. Of the synthetic repeats, 5 were chosen that best resemble bell-shaped distributions for the target and control values.

The following applies to each of the synthetic repeats. Four noise levels were used: *g* = 0.1, 0.2, 0.4, 0.8 (**Fig. 2**). The noise is applied as the product of three components: the noise level *g*, the standard deviation of the original outputs *σ_original_*, and samples from a truncated normal distribution. The resulting noisy output values are defined by the following equation:

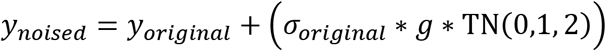

where:

*y_original_* = original output values
*σ_original_* = standard deviation of the original output
*g* = noise level
TN(0,1, 2) = a sample from a truncated normal distribution with mean 0, standard deviation 1, and bounds of ±2 standard deviations.

This application of noise ensures that noise is scaled appropriately and bound to avoid extreme perturbations.

**Fig. 2.**
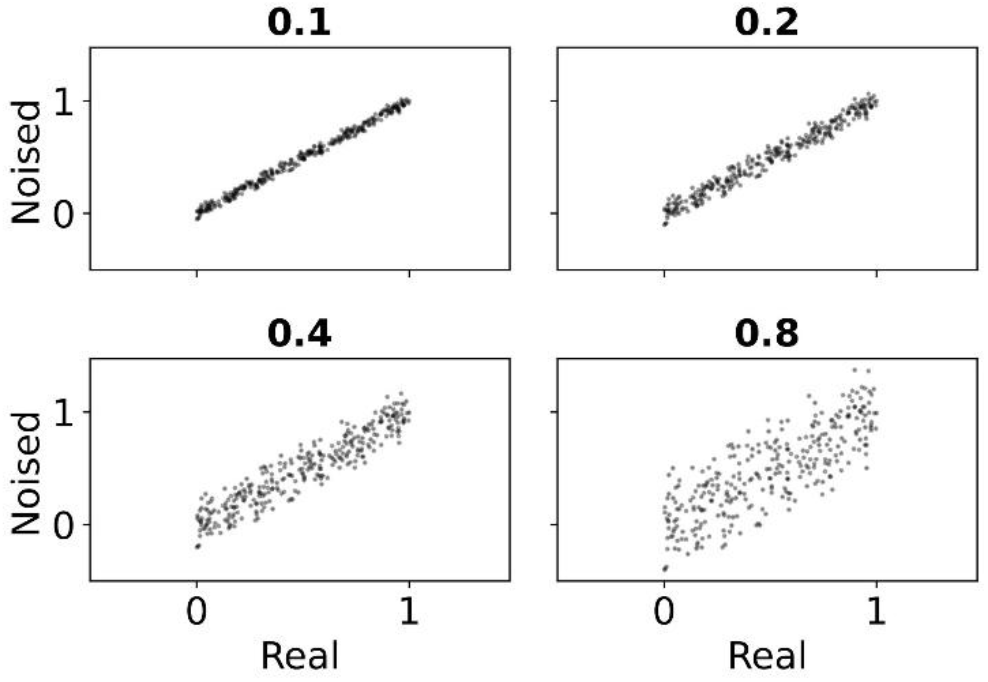
Examples of levels of noise. Different levels of noise were added (“Noised”) to a random sample of 300 values from a uniform distribution.

##### 3.1.2.1 Right-Skew

The following applies to each level of noise. To facilitate downstream selection, only noised output values within two standard deviations were kept. The output values were then min-max scaled, yielding values that range from [0, 1]. Each value was assigned a weight by histogram binning with 1000 bins, determining the proportion of observations of each bin, and assigning each value the inverse of the proportion of their bin. Each value was then partitioned into 20 histogram bins.

Different levels of right-skew were determined as different *r* (-2, -4, -8) values of a decaying exponential function (*e^rx^*) (**Fig. 3**). For each right skew level, the proportion of observations per bin for 20 bins was determined.

**Fig. 3.**
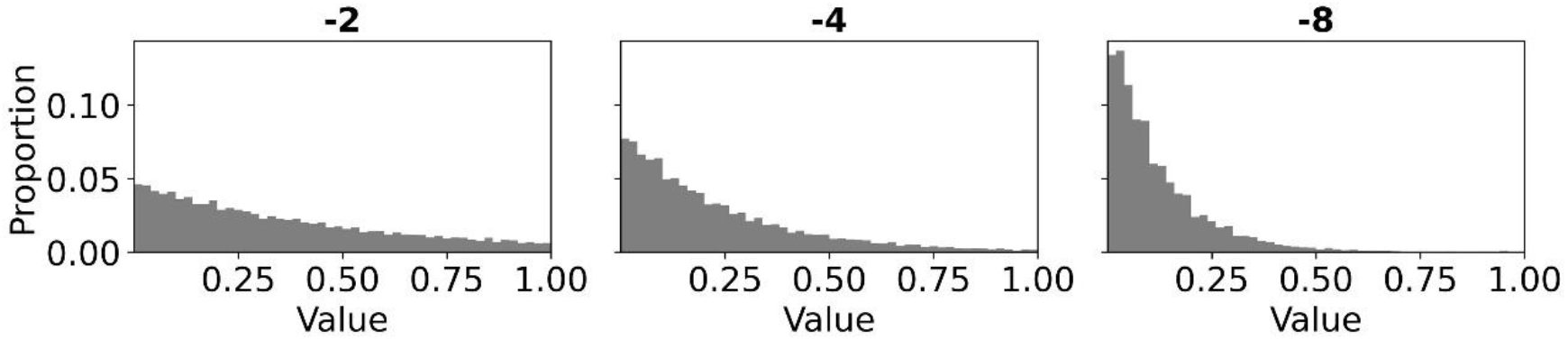
Examples of levels of right-skew. Histogram (50) bins of output values pertaining to different levels of right-skew.

##### 3.1.2.2 Number of Observations

For each combination of noise and right-skew, specific target counts were selected: *N* = 1000,2000,4000,8000, 16000. The required number of observations for any given bin *i* (*n_i_*) was determined by taking the product of the total desired observations *N* and the right-skew proportion for that specific bin (*P_i_*):

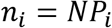

To populate each bin, *n_i_* observations were selected using random weighted sampling. This selection process used the inverse-proportion weights assigned to each observation earlier, ensuring intra-bin balance among the sampled data.

##### 3.1.2.3 Binning

For each dataset, the observations were binned using Minimum Transformation with Tanh as the transformation function. The number of bins and their ranges were determined with the following constraints: a minimum of 10 bins, a maximum of 50 bins, a minimum of 10 observations per bin, a scale factor minimum of 0.1, and a scale factor maximum of 3.

##### 3.1.2.4 Balanced Set

For downstream evaluation*, a balanced set* with a uniform distribution was selected from the unselected observations. The unselected observations were reweighted using the previous weighting scheme and 2000 observations were selected by random weighted sampling.

Each combination of synthetic repeat, noise, right-skew, and number of observations was repeated 5 times, referred to as processing repeats.

#### 3.1.3 Data Splits

The following applies to each combination of synthetic repeat, processing repeat, noise, right-skew, and number of observations. All splits had three divisions (training, validation, testing) with proportions of 0.6, 0.2, and 0.2 following popular data splitting regimes for modelling with NNs. For objective splitting, 1000 random splits were generated. For each split, for each division, the number of observations belonging to each bin were determined. The random splits with a division that contained 0 observations belonging to a bin were filtered out to facilitate later scoring.

A total of 12 metrics were used: random scoring, Wasserstein distance, KL divergence, reverse (also known as swapped) KL divergence, symmetric KL divergence, Shannon divergence, Pearson chi-squared, Neyman chi-squared, symmetric (also known as triangular) chi-squared, absolute relative difference (absrel), swapped absrel, and symmetric absrel. Using random scoring for objective splitting is equivalent to random split. All-against-one comparisons used all 12 metrics while All-against-all comparisons used only symmetric metrics: Wasserstein distance, symmetric KL divergence, Shannon divergence, symmetric chi-squared, and symmetric absrel.

Each of the filtered random splits were scored using all-against-one or all-against-all scoring. For each scoring method, the best split was kept. Generating, scoring, and selecting splits was repeated 20 times, referred to as split repeats.

#### 3.1.4 Model Fitting

All modelling used the Base NN, a batch size of 512, and a patience of 10. The choice of patience was informed by preliminary trials (not included here) which indicated that increasing patience did not yield significant performance improvements and this setting reduced computational overhead. Also, all models used the Adam optimizer (Kingma & Ba, 2014), selected for its widespread adoption and proven effectiveness across a broad range of NN applications. An Adam learning rate of 0.01 was used for all experiments with synthetic data, determined as the highest learning rate that yielded the greatest performance of learning rates that ranged from 0.0001 to 0.1 (data not included here).

Unless otherwise specified, the recip-MSE was used as the loss function and stopping criterion as it was shown to outperform other loss functions. For any MSE-related metric, lower scores are preferred.

For models fit with Epoch sampling, the weights for comp-MSE or recip-MSE are determined after sampling.

#### 3.1.5 Datasets Selection

For each combination of (noise level, right-skew level, number of observations), the predictions on the testing set across synthetic repeats, processing repeats, and split repeats were gathered and combined. Scatterplots were used to visualize the predictions versus actuals and combinations of (noise level, right-skew level) were chosen based on how well the models performed with the maximum number of observations. Combinations with noise levels 0.2 or 0.4 and right-skew levels of -2, -4, or -8, were chosen. For each selected (noise level, right-skew level), the number of observations was chosen as the minimum number of observations to achieve adequate performance. For combinations with right-skew level of -2, -4, and -8, the number of observations required was 1000, 2000, and 4000, respectively. Data is not included in this study but is present in the code repository.

#### 3.1.6 Model Scoring

For each modelling subject, modelling was repeated 15 times, generating 15 models. Bootstrapping simulated model repeats. For each set of bootstrapped repeats, the predictions of the top 3 scoring models (evaluated on the validation set) were averaged, akin to an average ensemble.

##### 3.1.6.1 Data Split Experiments

The average ensemble predictions on the testing set were scored using recip-MSE, referred to as bootstrapped scores. Across split repeats, the sets of bootstrapped scores were combined and summarized as median + median absolute deviation, referred to as the split score, to account for both the magnitudes and consistencies of the performances.

For every combination of (synthetic repeat, processing repeat, noise level, right-skew level), the split scores of the data split methods were ranked. The ranked split scores across synth repeats and processing repeats were combined and the mean rank was determined. For each combination of (noise level, right-skew), the rankings of the split methods were compared to the rankings of random split (which served as a baseline) by a one-sided ranked Wilcoxon signed-rank tests with *p*-value thresholds of 0.05.

Across all repeats and data combinations, the ranked scores were combined, and the mean rank was determined, yielding *overall* rankings. The rankings of the split methods were compared to the rankings of random split (which served as a baseline) and to the rankings of stratified split (which was the overall best scoring split method) through one-sided ranked Wilcoxon signed-rank tests with *p*-value thresholds of 0.05.

##### 3.1.6.2 Model Fitting Experiments

The average ensemble predictions on the balanced set were scored using Root Mean Squared Error (RMSE), referred to as bootstrapped scores. A subject model fitting method was compared to a reference model fitting method by the pairwise percent relative difference of corresponding bootstrapped scores, referred to as pairwise bootstrapped scores. Across synthetic repeats, processing repeats, and split repeats, the sets of pairwise bootstrapped scores were combined. The median was determined. For statistical assessment, the 5^th^ (*P*_5_) and the 95^th^ (*P*_95_) percentiles of the pairwise bootstrapped scores were determined, simulating a one-sided statistical significance level of 0.05 and a two-sided statistical significance level of 0.1 (Efron & Tibshirani, 1986). If 0 > *P*_95_, percentile, the subject model is significantly better than the reference model. If 0 < *P*_5_ percentile, the subject model is significantly worse than the reference model. If *P*_5_ < 0 < *P*_95_, the subject model is not significantly different from the reference model.

For experiments involving sampling, the ensembles were also scored by the resources used. In each epoch, the model is exposed to the training set data for fitting and the validation set data for scoring. When using sampling, a sample of the training set is used for fitting, but all the validation set is used for scoring. The resources used are calculated as:

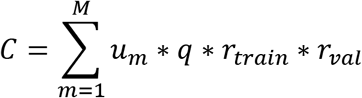

where:

*C* = resource cost
*M* = number of models
*u_m_* = number of epochs used by model *m* of *M*
*q* = sampling proportion
*r_train_*, *r_val_* = proportion of observations in the training and validation sets A lower resource cost indicates higher efficiency which is preferred.

### 3.2 Results and Discussion

#### 3.2.1 All-Against-One Objective Data Splitting with Absolute Relative Difference Yields Model Performances Similar to Stratified Split

Stratified random splitting yielded the best and most consistent model performances, as expected (**Table 1**). Of the objective data splitting methods evaluated, only all-against-one data splitting with absrel as the scoring function was significantly better than random data splitting (**Sup. Table 1**). It was also the only objective data splitting method that was not significantly worse than stratified random splitting. Thus, it is recommended that all-against-one objective data splitting with absrel be used for splitting non-independent imbalanced regression data. To the best of our knowledge, objective data splitting with absrel scoring has not been done before.

**Table 1.** Only stratified data splitting and all-against-one data splitting with absrel are significantly better than random data splitting. Data splitting methods were tried on datasets with different combinations of right-skew and noise. The mean rankings are presented; lower values are preferred. Bolded values in the overall rankings indicate methods that are significantly better than random data splitting. Full statistics in **Sup. Table 1**.

|  | Right-skew | -2 |  | -4 |  | -8 |  | Overall |
| --- | --- | --- | --- | --- | --- | --- | --- | --- |
|  | Noise Level | 0.2 | 0.4 | 0.2 | 0.4 | 0.2 | 0.4 |  |
| Stratified Splitting Methods |  |  |  |  |  |  |  |  |
|  | Stratified-Det | 5.7 | 7.9 | 7.4 | 7.0 | 8.0 | 9.1 | 7.5 |
|  | Stratified-Rand | 6.3 | 7.9 | 8.7 | 7.4 | 9.8 | 8.2 | 8.1 |
| Objective Splitting Methods |  |  |  |  |  |  |  |  |
| Against | Metric |  |  |  |  |  |  |  |
| One | Absrel | 8.1 | 7.8 | 7.7 | 7.6 | 8.2 | 7.4 | 7.8 |
| One | Wasserstein | 9.1 | 8.1 | 9.1 | 9.1 | 8.7 | 8.5 | 8.8 |
| One | Reverse KL | 10.1 | 8.2 | 9.4 | 9.8 | 7.8 | 7.6 | 8.8 |
| One | Neyman Chi-squared | 8.3 | 8.8 | 9.7 | 9.8 | 7.9 | 8.7 | 8.9 |
| One | Reverse Absrel | 10.5 | 8.5 | 8.8 | 7.0 | 9.7 | 8.8 | 8.9 |
| One | Sym. Absrel | 10.2 | 8.6 | 7.4 | 8.4 | 10.0 | 9.6 | 9.0 |
| All | Sym. Absrel | 8.5 | 10.7 | 9.8 | 8.4 | 9.6 | 7.8 | 9.1 |
| All | Sym. Chi-squared | 9.9 | 7.9 | 9.1 | 9.0 | 9.0 | 10.0 | 9.2 |
| One | Sym. KL | 9.7 | 9.3 | 10.6 | 8.1 | 7.9 | 9.5 | 9.2 |
| All | Wasserstein | 8.5 | 9.3 | 8.6 | 11.2 | 8.3 | 9.4 | 9.2 |
| All | Sym. KL | 10.7 | 9.6 | 8.8 | 8.6 | 8.2 | 9.4 | 9.2 |
| One | Random | 10.4 | 9.1 | 7.6 | 11.2 | 9.4 | 9.0 | 9.5 |
| One | Sym. Chi-squared | 8.4 | 8.6 | 9.6 | 9.3 | 11.1 | 10.8 | 9.6 |
| One | Pearson Chi-squared | 9.2 | 10.9 | 9.7 | 10.0 | 9.2 | 9.5 | 9.7 |
| One | KL | 9.2 | 11.8 | 11.1 | 11.0 | 10.2 | 9.6 | 10.5 |

#### 3.2.2 Models Fit with Recip-MSE Outperform Models Fit with Comp-MSE on Synthetically Generated Data

Models fit with comp-MSE as the loss function performed statistically similar to models fit with MSE as the loss function across all datasets (**Table 2**). Models fit with recip-MSE as the loss function performed better (if not similarly) than models fit with MSE as the loss function across all datasets and were statistically better on datasets with greater levels of right-skew and noise. Therefore, for modelling imbalanced regression, it is recommended that models are fit with recip-MSE as the loss function. Modelling with recip-MSE as the loss function produced well-performing model across all datasets (**Sup. Fig. 1**).

**Table 2.** Models fit with recip-MSE perform significantly better than models fit with MSE on synthetically generated data. Median pairwise bootstrap scores of models fit with different loss functions compared to models fit with MSE. Full statistics in Sup. Table 2. Lower values are preferred. Bolded values indicate that the comparator model performance is significantly different to the reference model performance.

| **Right-skew** | **-2** |  | **-4** |  | **-8** |  |
| --- | --- | --- | --- | --- | --- | --- |
| **Noise Level** | **0.2** | **0.4** | **0.2** | **0.4** | **0.2** | **0.4** |
| **Loss Func.** |  |  |  |  |  |  |
| MSE | 0.0 | 0.0 | 0.0 | 0.0 | 0.0 | 0.0 |
| comp-MSE | 0.2 | 0.0 | -0.9 | -0.8 | -1.5 | 0.7 |
| recip-MSE | -0.7 | -4.8 | -3.4 | **-13.8** | -4.5 | **-27.2** |

#### 3.2.3 Models Fit with Sampling Perform Worse than Models Fit with All Data but use Fewer Resources on synthetically generated data

Fitting models with recip-MSE and different sampling proportions was tried. Similar results are observed for sampling without replacement (**Table 3**, **Table 4**) and sampling with replacement (**Sup. Table 5, Sup. Table 7**). For brevity, only results of sampling without replacement are specifically referenced. Across all datasets, models fit with lower sampling proportions are progressively worse than models fit without sampling. The effects of sampling on performance are dampened with increasing right-skew and noise. On the datasets with the greatest right-skew and noise, models fit with sampling yield statistically similar performance to models fit without sampling, even with a sampling proportion of 0.05 (i.e. fitting with 5% of the training data). Furthermore, across all datasets, models fit with lower sampling proportions use increasingly less resources than models fit without sampling. Models fit with sampling proportions of 0.2 or less used significantly less resources than models fit without sampling.

**Table 3.** Models fit with sampling without replacement perform as well if not worse than models fit without sampling on synthetically generated datasets. Median pairwise bootstrap scores of models fit with different sampling proportions (“Samp. Prop.”) and without replacement compared to models fit without sampling (“None”). Full statistics in Sup. Table 3. See Table 2 for additional descriptions.

| **Right-skew** | **-2** |  | **-4** |  | **-8** |  |
| --- | --- | --- | --- | --- | --- | --- |
| **Noise Level** | **0.2** | **0.4** | **0.2** | **0.4** | **0.2** | **0.4** |
| **Samp. Prop.** |  |  |  |  |  |  |
| None | 0.0 | 0.0 | 0.0 | 0.0 | 0.0 | 0.0 |
| 0.8 | 4.7 | 2.2 | 1.1 | 0.2 | 0.5 | -0.6 |
| 0.6 | 7.5 | 3.0 | 1.8 | 0.5 | 1.6 | -0.7 |
| 0.4 | **10.7** | 4.1 | 8.8 | 2.7 | 2.9 | -0.4 |
| 0.2 | **17.0** | **6.2** | **13.7** | 4.0 | 10.5 | 1.1 |
| 0.1 | **40.2** | **14.1** | **21.3** | 5.8 | 11.2 | 1.2 |
| 0.05 | **83.9** | **46.8** | **60.6** | **37.2** | 13.8 | 2.3 |

**Table 4.**
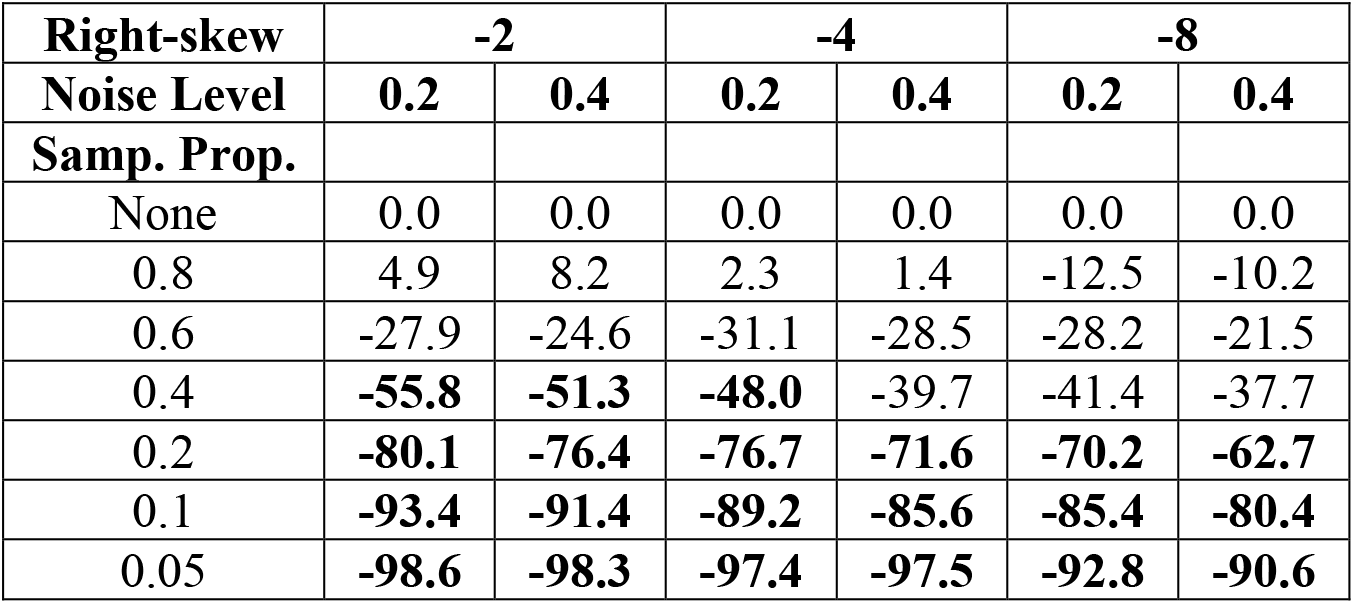
Models fit with sampling without replacement and proportions of 0.2 or less used significantly fewer computing resources than models fit without sampling on synthetically generated datasets. Median resource pairwise bootstrap scores of models fit with different sampling proportions (“Samp. Prop.”) and without replacement compared to models fit without sampling (“None”). Full statistics in **Sup. Table 4**. See Table 2 for additional descriptions.

Although fitting with sampling does not improve model performance and may yield lower model performance, the monotonic relationships of model performance and computational cost to sampling proportions are useful for real-world data settings. In this case study, the same NN architectures were used to produce the data and model the data. Hence, it was assumed that the configuration of the non-learnable parameters was optimal for modelling the data. For real-world data, however, the optimal configuration of the non-learnable parameters of a model is not known.

Different configurations are tried toward determining the optimal configuration in a process known as hyperparameter optimization (HPO). A greater number of configurations tried may yield better models and thus better final performance, but each additional configuration tried increases the computational cost. Real-world data, especially genomics data, is often large and complex, requiring larger and more complex models. When using the entire data, only relatively few configurations can be tried.

Assuming that the performances of configurations under sampling are related to the performances when using the full data, sampling can be used in HPO to reduce the computational cost of each configuration tried. This enables more configurations to be tried with the same computational cost, ultimately producing better performing models. It is recommended that HPO yield multiple configurations and a final test involving the full data is tried to better ensure correct selection.

## 4 Case Study: Real Genomics Datasets

This case study presents examples of two popular genomics modelling tasks: protein-DNA interactions and enhancer activity. Both tasks are single-output regression tasks, and the outputs are right-skewed. Different model fitting methods are tried toward improving model performance.

The interactions of proteins with genome DNA are the principal components of gene regulation. These interactions are influenced by the DNA sequence and/or surrounding environment (e.g. histone post-translational modifications) (Guertin & Lis, 2010; Slattery et al., 2014; Wang et al., 2012). Yin Yang 1 (YY1) is a DNA sequence-specific transcription factor (a protein that affects gene transcription) implicated in human diseases such as cancer and has a significant role in development, specifically neurogenesis (Verheul et al., 2020). CTCF is also a DNA sequence-specific transcription factor influential in chromatin architecture (Phillips & Corces, 2009). The protein-genome interactions of YY1 and of CTCF in a human cancer cell line are modelled here.

DNA sequences possess different abilities to regulate gene expression. The ability of a specific DNA sequence to positively influence gene regulation is known as enhancer activity. The enhancer activities of DNA sequences in a human cancer cell line are modelled here.

For each task, three different NN models are used. For modelling protein-DNA interactions, Bichrom (Srivastava et al., 2021), Multi-Mimic (Abdulnabi & Westwood, 2026a), and Multi-Oyster (Abdulnabi & Westwood, 2026b) are used. For modelling enhancer activities, DeepSTARR (de Almeida et al., 2022), Mimic (Abdulnabi & Westwood, 2026a), and Oyster (Abdulnabi & Westwood, 2026b) are used. Together, model fitting methods are experimented on 3 real genomics datasets with an assortment of different models.

### 4.1 Methods

See https://github.com/Husam94/CS-YY1/, https://github.com/Husam94/CS-CTCF/, https://github.com/Husam94/CS-EnhAct/ for full code on YY1-DNA, CTCF-DNA, and enhancer activity experiments, respectively. Descriptions on data processing and model configurations are also provided in Abdulnabi & Westwood (2026a). Only relevant information is included here.

**Fig. 4.**
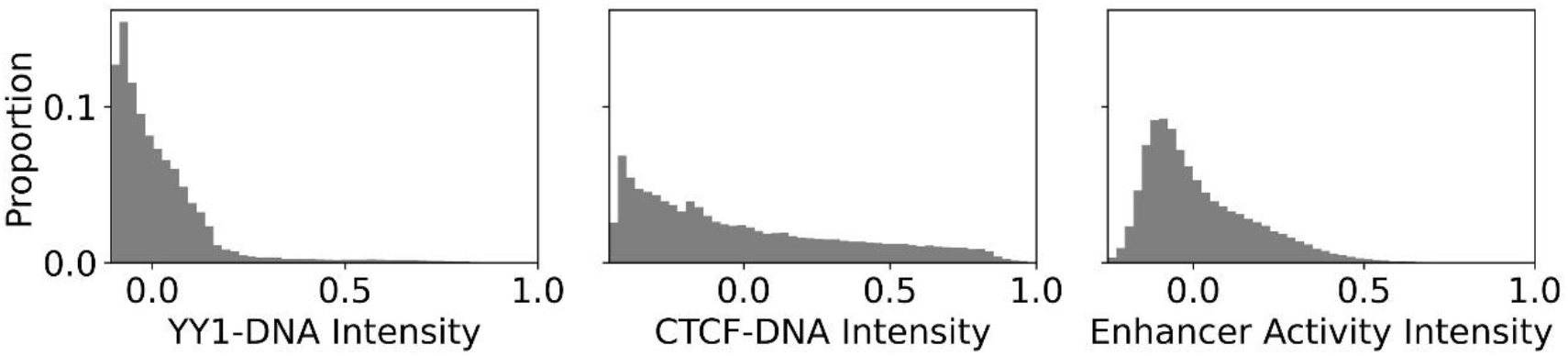
Output values of genomics datasets are right-skewed. Histograms (50 bins) of YY1-DNA intensity (left), CTCF-DNA intensity (right), and enhancer activity intensity (right).

#### 4.1.1 Binning and Data Splitting

Refer to **Section 3.1.2.3** for binning details.

Datasets were split into 4 sets with proportions of 0.55, 0.15, 0.15, and 0.15. The first 3 sets are the training, validation, and testing. The last set is an additional holdout set to evaluate the final selected model in the case study that follows in Abdulnabi & Westwood (2026b). The datasets were split using objective data splitting with absrel scoring and 100,000 random splits. Observations within 50,000 nucleotides on the same chromosome were grouped together prior to splitting.

#### 4.1.2 Model Scoring

Model Scoring is the same as in **Section 3.1.6** except that the models were scored on the testing set – there is no balanced set for either task. The testing set is imbalanced, so the reciprocal density weights were used in RMSE (recip-RMSE).

#### 4.1.3 Model Configurations and Model Fitting

For each dataset, for each type of model, the configurations of non-learnable tunable parameters were determined by a two-step procedure. First, 300 random configurations were generated and fit using recip-MSE as the loss function and Epoch Sampling with a proportion of 0.1 and the top 5 configurations were selected. The configurations included batch size, optimizer, and learning rate. All models were fit using a patience of 10. Second, each of the top 5 configurations were fit with recip-MSE as the loss function but without Epoch Sampling and repeated 10 times. Each configuration was scored with mean bootstrap scores, and the top scoring candidate was chosen for further experiments.

### 4.2 Results and Discussion

Models fit with comp-MSE as the loss function performed similar to models fit with MSE as the loss function across all models and datasets except for two instances: Bichrom models on YY1-DNA data performed statistically better by 14.2% and Multi-Mimic models on CTCF-DNA data performed statistically worse by 2.7% (**Table 5**). Models fit with recip-MSE as the loss function performed statistically significantly and substantially better than models fit with MSE as the loss function across all models and datasets by 7.8-40.9%. Modelling with recip-MSE as the loss function yielded well performing models across all datasets (**Sup. Fig. 2**).

**Table 5.**
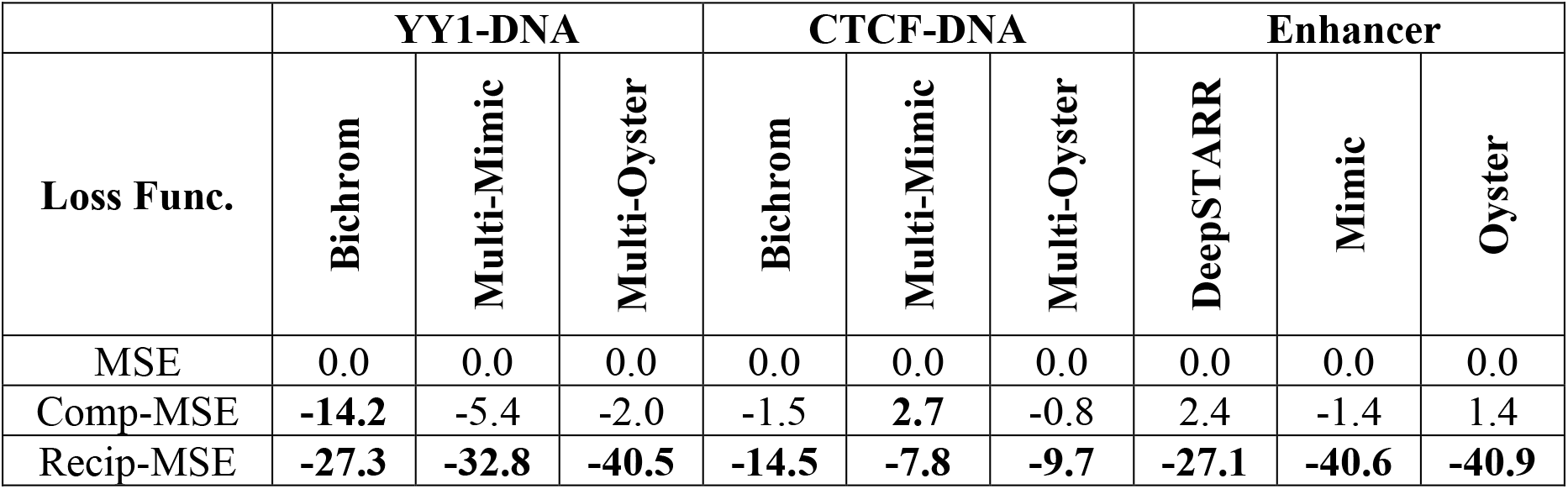
Models fit with recip-MSE perform significantly better than models fit with MSE on real genomics datasets. Median pairwise bootstrap scores of models fit with different loss functions compared to models fit with MSE. Full statistics in **Sup. Table 9**. See Table 2 for additional descriptions.

Fitting models with recip-MSE and different sampling proportions without replacement was tried (**Table 6**). Models fit with different sampling proportions without replacement performed similarly to models fit without sampling across all models and datasets except for Multi-Oyster models on YY1-DNA and Bichrom models on CTCF-DNA that performed significantly worse by 4.0 and 3.5%, respectively, when fit with a sampling proportion of 0.05.

**Table 6.** Models fit with sampling without replacement generally perform similarly to models fit without sampling on real genomics data with exceptions. Median pairwise bootstrap scores of models fit with different sampling proportions (“Samp. Prop.”) and without replacement compared to models fit without sampling (“None”). Full statistics in **Sup. Table 10**. See Table 2 for additional descriptions.

|  | YY1-DNA |  |  | CTCF-DNA |  |  | Enhancer |  |  |
| --- | --- | --- | --- | --- | --- | --- | --- | --- | --- |
| Samp. Prop. | Bichrom | Multi-Mimic | Multi-Oyster | Bichrom | Multi-Mimic | Multi-Oyster | DeepSTARR | Mimic | Oyster |
| None | 0.0 | 0.0 | 0.0 | 0.0 | 0.0 | 0.0 | 0.0 | 0.0 | 0.0 |
| 0.8 | 5.7 | -1.8 | -0.5 | 0.2 | 0.6 | -0.5 | -1.4 | -0.3 | 1.9 |
| 0.6 | 5.2 | 0.1 | -2.1 | 1.0 | 0.4 | -0.2 | -0.9 | 1.0 | 1.9 |
| 0.4 | 2.1 | 1.8 | -0.8 | 0.7 | 1.0 | -0.2 | 0.1 | -2.2 | 0.3 |
| 0.2 | 9.6 | 2.7 | -2.8 | 2.6 | 0.8 | -0.2 | 1.1 | -2.9 | 0.5 |
| 0.1 | 7.9 | 5.1 | 0.3 | 2.4 | 1.0 | 0.2 | 7.9 | -2.2 | 1.6 |
| 0.05 | 6.1 | 0.7 | **4.0** | **3.5** | 2.4 | 0.6 | -0.3 | -1.7 | 0.0 |

Fitting models with recip-MSE and different sampling proportions with replacement was tried (**Table 7**). Models fit with different sampling proportions with replacement generally performed similarly to models fit without sampling, but a few exceptions led to significantly worse performance by 2.2 to 10.5%.

**Table 7.** Models fit with sampling with replacement generally perform similarly to models fit without sampling on real genomics data with exceptions. Median pairwise bootstrap scores of models fit with different sampling proportions (“Samp. Prop.”) and with replacement compared to models fit without sampling (“None”). Full statistics in **Sup. Table 11**. See Table 2 for additional descriptions.

| Samp. Prop. | YY1-DNA |  |  | CTCF-DNA |  |  | Enhancer |  |  |
| --- | --- | --- | --- | --- | --- | --- | --- | --- | --- |
|  | Bichrom | Multi-Mimic | Multi-Oyster | Bichrom | Multi-Mimic | Multi-Oyster | DeepSTARR | Mimic | Oyster |
| None | 0.0 | 0.0 | 0.0 | 0.0 | 0.0 | 0.0 | 0.0 | 0.0 | 0.0 |
| 1 | 4.0 | 3.9 | **8.5** | 1.5 | 2.3 | 0.1 | 6.8 | 2.6 | **3.7** |
| 0.8 | 8.0 | 2.8 | 0.1 | 0.0 | 1.7 | 0.1 | 1.7 | 1.2 | **3.0** |
| 0.6 | -0.6 | 5.4 | **4.0** | -0.6 | 0.9 | -0.1 | 0.9 | **6.6** | 2.0 |
| 0.4 | 16.4 | 1.9 | **5.2** | 0.9 | 2.0 | -0.3 | -1.2 | **7.3** | **2.2** |
| 0.2 | 3.7 | -1.4 | 3.5 | 0.2 | 1.0 | -0.2 | 2.1 | **7.8** | 1.1 |
| 0.1 | 15.9 | 2.9 | 2.5 | 2.1 | 1.5 | 0.0 | 4.1 | **10.5** | 2.2 |
| 0.05 | 15.5 | 1.7 | 2.5 | **3.8** | 2.4 | 0.8 | **9.0** | **10.5** | **2.8** |

In general, across all datasets and model configurations, sampling with or without replacement did not yield better performing models. While models fit with sampling used substantially and significantly less resources than models fit without sampling (data not included here but see the accompanying code), this advantage becomes irrelevant after HPO.

The configurations of all models in this case study were determined through a two-stage HPO based on the prescription of the previous case study: a random search with models fit with recip-MSE as the loss function and sampling followed by repeat fitting of top configurations without sampling. After HPO, the objective shifts entirely to maximizing predictive performance; computational savings from sampling are no longer relevant.

Based on these findings, it is recommended that recip-MSE is used as the loss function without sampling for modelling genomics data. Still, this case study is limited in scope; it remains possible that fitting with sampling may improve model performance of other models on other datasets.

## 5 Conclusion

Genomics data is often right-skewed and involves non-independent observations which complicates model scoring, data splitting, and model fitting, thereby requiring special care at each of these steps. Binning is an effective method for density estimation and facilitates model scoring and data splitting. Minimum Transformation Binning is a novel objective binning method that improves density estimation. Stratified splitting provides the optimal partition for independent observations but cannot be used for non-independent observations. Objective data splitting improves on Random split for splitting non-independent observations, but which internal scoring function to use is not obvious. Through experiments with synthetically generated data, it was found that the absrel scoring function is the most effective out of 11 scoring functions tried—a scoring function for data splitting that has not been used before, to the best of our knowledge. Across synthetic and real-world genomics datasets, model fitting with recip-MSE consistently yielded statistically significant performance gains and thus is recommended for model fitting imbalanced data. Model fitting with sampling, however, generally did not improve model performance but may use substantially less resources and may be useful for HPO. Together, this study provides a unified framework for modelling imbalanced regression: bin and estimate density by Minimum Transformation Binning; split non-independent data with absrel objective data splitting; and score and fit models with recip-MSE.

## 6 Acknowledgements

We thank Anthony Bonner and Alan Moses from the University of Toronto for constructive discussions.

## 7 Author Contributions

**Conceptualization**: Husam Abdulnabi

**Data curation**: Husam Abdulnabi

**Formal analysis**: Husam Abdulnabi

**Investigation**: Husam Abdulnabi

**Methodology**: Husam Abdulnabi

**Project administration:** Husam Abdulnabi

**Software**: Husam Abdulnabi

**Supervision**: J. Timothy Westwood

**Writing – original draft:** Husam Abdulnabi

**Writing – review & editing:** J. Timothy Westwood

## 9 Supplementary

### 9.1 Figures

**Sup. Fig. 1.**
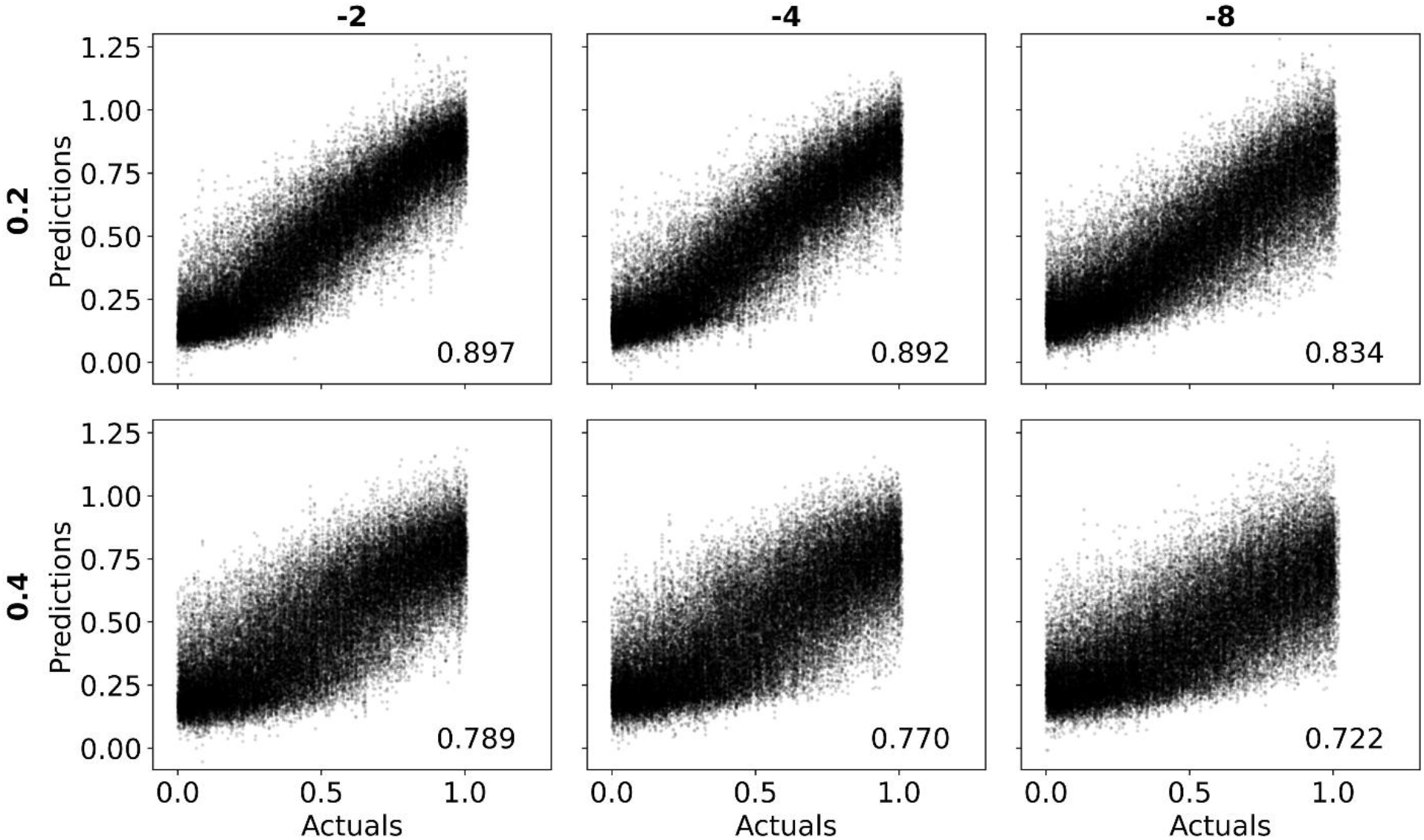
Models fit with recip-MSE as the loss function perform sufficiently well across all synthetically generated datasets. Scatterplots visualizing the average of the predictions of the bootstrap ensembles versus actuals of the balanced sets of datasets with different combinations of right-skew (per column) and noise levels (per row). Each plot depicts predictions versus actuals across all corresponding synthetic, processing, and split repeats and the recip-PCC is included at the bottom right.

**Sup. Fig. 2.**
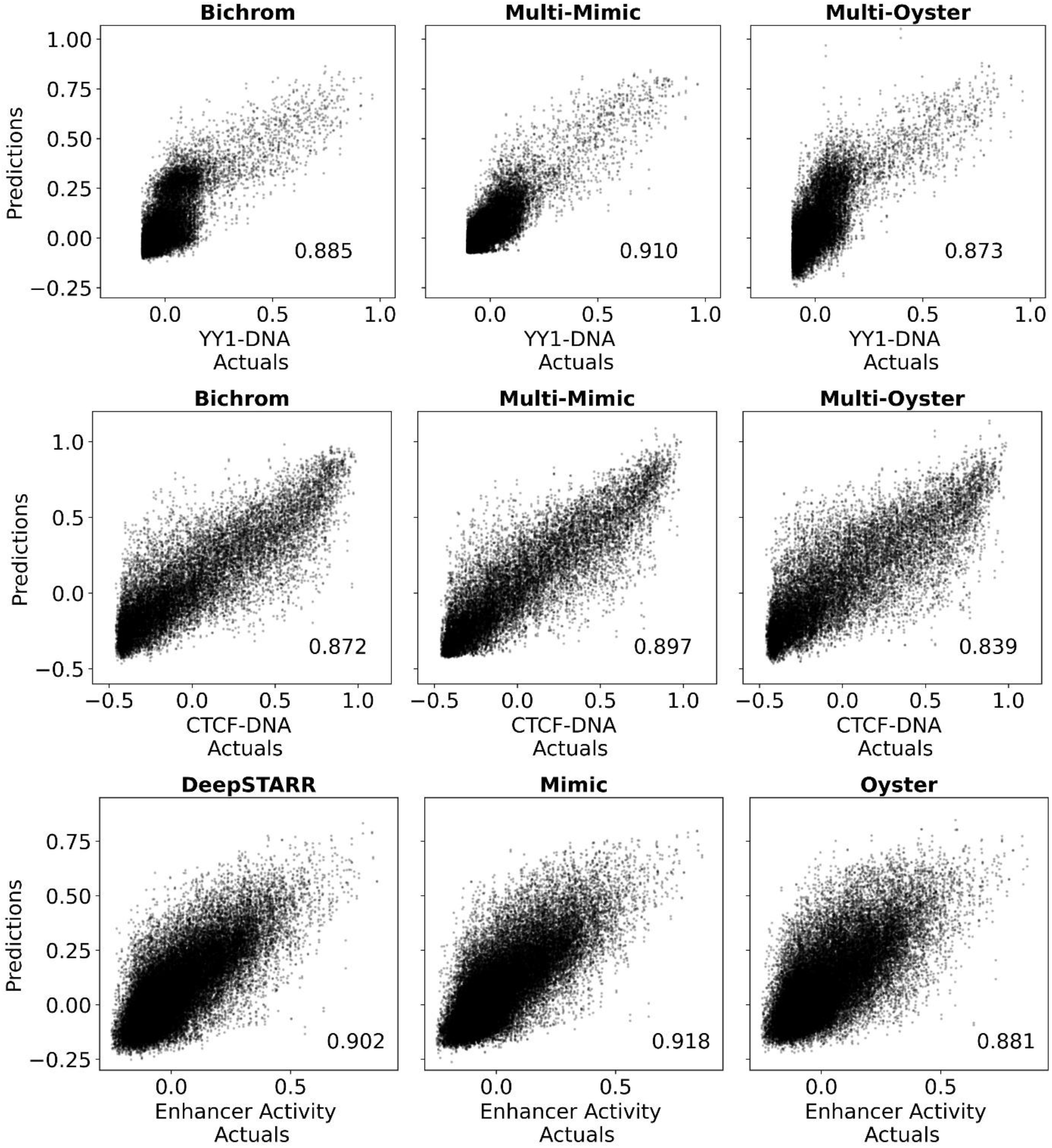
Models fit with recip-MSE as the loss function perform well on real genomics datasets. Scatterplots visualizing the average of the predictions of the bootstrap ensembles versus actuals of the testing set of Bichrom or DeepSTARR, Mimic, and Oyster models fit on YY1-DNA interactions data (first row), CTCF-DNA interactions data (second row) and on enhancer activity data (third row). The recip-PCC is included at the bottom right of each plot

### 9.2 Tables

All tables in BinMeths.Sup_Tables.xlsx

#### 9.2.1 Synthetic data case study

**Sup. Table 1.** Statistics comparing data split methods to random split and to stratified random split on synthetically generated datasets. Difference in rank to reference (“Rank Diff.”) is reported. Comparing to random split, Wilcoxon *p*-value of better than random split is reported. Compared to stratified random split, Wilcoxon *p*-value of worse than stratified random split is reported. *p*-values are bolded if less than 0.05.

**Sup. Table 2.** Statistics of pairwise bootstrap scores comparing models with different loss functions to models fit with MSE as the loss function on synthetically generated datasets. Mean and median are reported as well as the 5^th^ (“P5”) and 95^th^ (“P95”) percentiles and whether the comparison is statistically significant (“Signif.”).

**Sup. Table 3.** Statistics of pairwise bootstrap scores comparing models fit with different sampling proportions without replacement to models fit without sampling on synthetically generated datasets.

**See Sup. Table 2** for additional descriptions.

**Sup. Table 4.** Statistics of resource pairwise bootstrap scores comparing models fit with different sampling proportions without replacement to models fit without sampling on synthetically generated datasets.

**See Sup. Table 2** for additional descriptions.

**Sup. Table 5.** Median pairwise bootstrap scores of models fit with different sampling proportions (“Samp. Prop.”) with replacement compared to models fit without sampling (“1”) on synthetically generated datasets. Full statistics in **Sup. Table 6.**

**Sup. Table 6.** Statistics of pairwise bootstrap scores comparing models fit with different sampling proportions with replacement to models fit without sampling on synthetically generated datasets.

See **Sup. Table 2** for additional descriptions.

**Sup. Table 7.** Median resource pairwise bootstrap scores of models fit with different sampling proportions (“Samp. Prop.”) with replacement compared to models fit without sampling (“1”) on synthetically generated datasets. Full statistics in **Sup. Table 8.**

**Sup. Table 8.** Statistics of resource pairwise bootstrap scores comparing models fit with different sampling proportions with replacement to models fit without sampling on synthetically generated datasets.

See **Sup. Table 2** for additional descriptions.

#### 9.2.2 Real Genomics Datasets Case Study

**Sup. Table 9.** Statistics of pairwise bootstrap scores comparing models with different loss functions to models fit with MSE as the loss function on real genomics datasets.

See **Sup. Table 2** for additional descriptions.

**Sup. Table 10.** Statistics of pairwise bootstrap scores comparing models fit with different sampling proportions without replacement to models fit without sampling on real genomics datasets.

See **Sup. Table 2** for additional descriptions.

**Sup. Table 11.** Statistics of pairwise bootstrap scores comparing models fit with different sampling proportions with replacement to models fit without sampling on real genomics datasets. See **Sup. Table 2** for additional descriptions.

## Notes

https://github.com/Husam94/poseigen_binmeths

https://github.com/Husam94/CS-Synth

https://github.com/Husam94/CS-YY1

https://github.com/Husam94/CS-CTCF

https://github.com/Husam94/CS-EnhAct

### Competing Interest Statement

The authors have declared no competing interest.

### Summary of Updates

Revised novelty claims, revised binning methods, new transformation binning method, more statistically robust and streamlined results, additional biological case studies.

